# Vitality capacity preservation through lifelong aerobic exercise: a pathway to healthy ageing

**DOI:** 10.64898/2026.06.04.728174

**Authors:** J. Mercier, O. Guérin, E. Michel, F. Chorin, A. Loubat, N. Gautier, A.S. Rousseau, S.S. Colson

## Abstract

**Background:** The distinction between healthy and pathological ageing has led to the concept of vitality capacity (VC), which can be understood as the body’s physiological reserve. An individual’s VC can be estimated using 12 biomarkers spread across 3 domains: immune and stress response, energy and metabolism and neuromuscular function. Vitality capacity may be preserved by lifelong physical activity. This cross-sectional study aimed to examine the relationship between lifelong aerobic physical activity and VC.

**Methods:** VC of 20 lifelong active and 19 inactive healthy adults aged >55 years was assessed using 12 biomarkers across the three VC domains. Domain-specific z-scores were calculated and averaged to derive a global VC score. Principal component analysis was performed and loadings extracted to estimate domains weight, and multiple correlations were conducted to identify associations among biomarkers, domains and VC scores.

**Results:** VC was higher in lifelong active participants (+0.2 z-score units, p = 0.006) and correlated with age (r = -0.53, p < 0.001). Neuromuscular domain contributed most to VC variability, and the immune and stress response domain was higher in the active group (+0.4 z-score units, p = 0.001) as energy/metabolism among female participants (+0.5 z-score units, p.adj = 0.006).

**Conclusion:** Lifelong aerobic physical activity is associated with higher VC in older adults, particularly within the immune and stress response domain. These findings highlight the role of physical activity in preserving the physiological reserve and reinforce the relevance of lifelong aerobic physical activity as a driver of healthy ageing.

## 1. Introduction

In 2015, the United Nations declared 2021-2030 the Decade of Healthy Ageing, promoting a shift from a disease-centered model of ageing toward the preservation of functional ability—the capacity of older adults to be and do the things they have reason to value^1^. This global paradigmatic shift has led to new definitions of healthy ageing and highlights the need for appropriate reference populations to study it.

Physically active older adults represent a particularly relevant model in this context, as the benefits of physical activity (PA) on health, ageing or longevity are well established and widely documented in numerous studies and systematic reviews^2–4^. Physically active older adults have been extensively studied and show increased longevity^5^, longer telomeres^6^ and improved inflammatory profile^7^, particularly in the context of aerobic training. While resistance training also provides age-related health benefits, particularly through improvements in body composition^8^, aerobic physical activity has the strongest associations with reduced mortality, cardiovascular protection, respiratory and metabolic health, and others broad systemic benefits^8–11^.

Beyond these traditional health outcomes, a recent study showed a positive association between moderate to vigorous physical activity (MVPA) and intrinsic capacity, a key concept in healthy ageing^12^. Intrinsic capacity is defined as the composite of all physical and mental capacities that an individual can draw on at any point in time that, together with environmental factors, would determine functional ability, and thus healthy ageing^13^. The physiological underlying domain of intrinsic capacity has been defined as vitality capacity (VC): a physiological state (due to normal or accelerated biological ageing processes) resulting from the interaction between multiple physiological systems, reflected in (the level of) immune and stress response, energy and metabolism, and neuromuscular function of the body^14^.

The potential contribution of physical activity (PA) to the preservation of VC has recently been reviewed^15^. Authors describe how PA, and particularly structured exercise, induces beneficial adaptations across the three domains of VC (i.e., immune and stress response, energy and metabolism, and neuromuscular function) thereby helping to maintain VC and support healthy ageing. Despite being reviewed, the relationship between PA and VC has not been demonstrated yet. Moreover, attempts to assess and/or establish VC score remain limited. A study reported positive associations between VC and cardiometabolic health and negative associations with various chronic diseases and impaired physical function^16^. However, the scoring system proposed^16^ was based on nine biomarkers among the 12 suggested by the WHO working group^14^. A recent study studied two domains of VC (i.e., energy and metabolism, and neuromuscular function) to validate their use in VC assessment and examined their links with locomotor function and quality of life^17^. However, as stated by the authors, the absence of immune and stress response domain highlights the need for further studies encompassing all three domains. These limitations are consistent with findings from a scoping review^18^, which highlighted a lack of alignment with the WHO framework and emphasized the need for standardized tools and approaches consistent with WHO recommendations.

Therefore, the present study operationalized a VC construct encompassing all three domains and aligned it with the 12 biomarkers proposed by the WHO working group^14^, by investigating the influence of lifelong aerobic PA practice on overall VC as well as the relative contribution of its domains using principal component analysis. We hypothesized that healthy older adults with a lifelong history of MVPA would exhibit higher VC than their inactive counterparts.

## 2. Method

### 2.1. Participants

This cross-sectional observational study included 40 healthy adults (20 males and 20 females) aged over 55 years, comprising 20 lifelong active and 20 lifelong inactive participants. All participants were free of chronic disease, non-smokers (or abstinent >3 years), and postmenopausal females had confirmed menopause for at least 6 months. Written informed consent was obtained from all participants, and the study was approved by the Comité de Protection des Personnes Ile de France II (approval no. 2024-A01310-47).

Active individuals were defined as those who had engaged in structured aerobic activity ≥3 times per week for the past 30 years, with no break longer than one week in the 3 months preceding the study and no break exceeding 12 consecutive months over that period. Their activity levels met or exceeded international recommendations for physical activity (WHO, 2020; ANSES, 2016), corresponding to ≥150 minutes per week of moderate-intensity aerobic activity, ≥75 minutes of vigorous activity, or an equivalent combination.

Inactive individuals were defined as those who did not meet these criteria, reporting predominantly sedentary or low-intensity occupational histories, no regular sports or fitness activity, and ≤75 minutes per week of leisure-time physical activity or transportation-related physical activity over the past 30 years.

Exclusion criteria included chronic disease, disability, hospitalization in the last 6 months, cardiovascular events, current medication, acute inflammation, history of doping, disability, legal guardianship. Participants were recruited in the Nice metropolitan area through associations, sporting events, local media, and social networks. Participants who reported a health issue during the experiment were removed from the study.

### 2.2. Procedure

The study comprised three visits. Visit 1 served as a screening session, whereas visits 2 and 3, conducted in the morning served as testing session. During visit 1, participants completed a semi-structured interview on physical activity history and the Global Physical Activity Questionnaire (GPAQ). Visit 2, conducted under fasting and resting conditions, included blood sampling, body composition assessment, and measurements of heart rate variability, oxygen saturation, and spirometry. Participants also completed dietary and self-reported questionnaires and were provided with a wrist accelerometer (worn for 7 days) and a food diary. During visit 3, participants returned the accelerometer and underwent, after a standardized warm-up, some physical function assessments, maximal handgrip strength, muscular endurance, and maximal isometric knee extensor strength.

### 2.3. Physical activity (PA) assessments

PA history was collected through a semi-structured interview conducted by an exercise science specialist, covering type, frequency, duration, and continuity of PA over the past 30 years, as well as occupational activity and active transportation. Current activity levels were further assessed using the Global Physical Activity Questionnaire (GPAQ^19^) with the assistance of the investigator. To obtain objective PA measures, participants wore a wrist-mounted accelerometer (AX3; Axivity, designed by Open Lab, Newcastle University) for 7 consecutive days. The AX3 accelerometer has been validated in adult and older populations^20,21^. Accelerometer data were analyzed using 1-second epochs and wrist-specific cut-points implemented in OmGui software (v1.0.0.45; Open Movement, Newcastle, UK). Participants were instructed not to remove the device and to maintain their usual PA habits during this period.

### 2.4. Quality of life assessment

The French version of the WHOQOL-OLD was used to assess quality of life in older adults^22^. This validated questionnaire comprises 24 items distributed across six domains: sensory abilities, autonomy, past–present–future activities, social participation, death and dying, and intimacy. Higher scores indicate better perceived quality of life.

### 2.5. Vitality Capacity assessments

#### 2.5.1. Immune and Stress Response

*IL-6 and TNF-α concentrations* (pg/mL) were quantified in plasma samples using the VPLEX multiplex immunoassay kit (MSD, Rockville, MD) according to the manufacturer’s instructions.

*Perceived immune status* (score) was measured using the Immune Status Questionnaire^23^. Oxygen saturation (%) was measured after 10 minutes of supine rest using a digital oximeter (CMS50D, Contec Medical Systems Co., Qinhuangdao, China).

*Autonomic function* (RMSSD, ms) was assessed via heart rate variability (HRV) measures. Short-term HRV was recorded with a chest strap sensor (Polar H10, Finland) and analyzed using Kubios software (Kuopio, Finland). Two 3-minute recordings were performed during the 10 minutes of supine rest. The root mean square of successive differences (RMSSD) was then extracted and reported.

#### 2.5.2. Energy and Metabolism

*Adiponectin* (µg/mL) was measured in plasma samples using the Human Adiponectin ELISA Kit (PR-KE00290-96T, Euromedex, Souffelweyersheim, France) following the manufacturer’s instructions.

*Glycated hemoglobin* (%) was measured in whole blood samples using Human Hemoglobin A1c Assay Kit (catalog #80099, Crystal Chem, Itaska, IL 60143 USA) following the manufacturer’s instructions.

*Body composition* (lean mass, %) was determined via dual-energy X-ray absorptiometry (Stratos DR, France).

*Nutritional status* (score) was assessed using the Mini Nutritional Assessment (MNA) questionnaire^24^.

*Perceived fatigue* (score) was evaluated using the French version of the Rating-of-Fatigue Scale questionnaire^25^.

*Muscular endurance* (time, s) was measured in the right hand and defined as the duration for which participants could maintain 50% of their maximal handgrip strength ^26^ on a manual dynamometer (K-grip, Kinvent, Montpellier, France) in a seated position with the elbow flexed at 90°. Verbal encouragement was provided.

#### 2.5.3. Neuromuscular Function

Knee extensor strength (N·m) was measured on right lower limb via maximal isometric contractions with knee bend at 90° on an ergometer (Biodex System 4 Pro, Biodex Medical, Shirley, New York, USA). Visual feedback and verbal encouragement were provided. Results were normalized to the right lower limb lean mass obtained from dual-energy X-ray absorptiometry testing.

*Respiratory muscle strength* was assessed through respiratory muscle function using the proportion of the person’s vital capacity that they are able to expire in the first second of forced expiration (FEV1/FVC) in a seated position using spirometry (microQuark, Cosmed, Italy), with participants performing a maximal inhalation and exhalation after 2 normal breaths. FEV1/FVC ratio was normalized using the deviation from the predicted score with the rspiro R package.

*Handgrip strength* (kg) was measured on right hand using a manual dynamometer (K-grip, Kinvent, Montpellier, France) during maximal voluntary contraction in a seated position with the elbow flexed at 90°. Result was normalized to the right upper limb lean mass obtained from dual-energy X-ray absorptiometry testing.

For all neuromuscular function biomarkers measurement, verbal encouragement was provided, and the highest value out of three trials was used for analysis.

### 2.6. Vitality Capacity score construct

To our knowledge, only one study proposed a VC score construct^16^. However, the unequal number of biomarkers across components (e.g., four energy and metabolism biomarkers, two neuromuscular function biomarkers) was not accounted, potentially biasing the final VC score towards energy and metabolism domain. We therefore propose a score construct that gives the same weight to each domain.

Vitality capacity, defined as a 12-biomarker construct^14^, was assessed by normalizing each individual biomarker into z-scores and aggregating them within their respective domain. For instance, neuromuscular function domain was calculated as the mean of z-scored handgrip strength, knee strength, and spirometry. For perceived fatigue, because a high fatigue z-score was associated with more fatigue, the score has been reversed to match with others. A same reverse logic has been applied to IL-6, TNF-α and HbA1c. As no specific blood biomarkers were predefined, four candidate biomarkers were measured, and those demonstrating the best fit with the data were retained for inclusion in the final construct. To this end, we compared all combinations of two inflammatory biomarkers (IL-6, TNF-α) and two metabolic biomarkers (adiponectin, HbA1c), making four different pairs of biomarkers, each one added to the remaining ten standardized vitality capacity biomarkers. Structural coherence was assessed using the variance explained by the first principal component (PC1), Cronbach’s alpha, and mean absolute inter-item correlation. IL-6 and adiponectin combination demonstrated greater dimensional dominance, internal consistency and comparable redundancy relative to the three others pairings (Fig. 1). Therefore, IL-6 and adiponectin has been retained and TNF-α and HbA1c removed from the VC score construct.

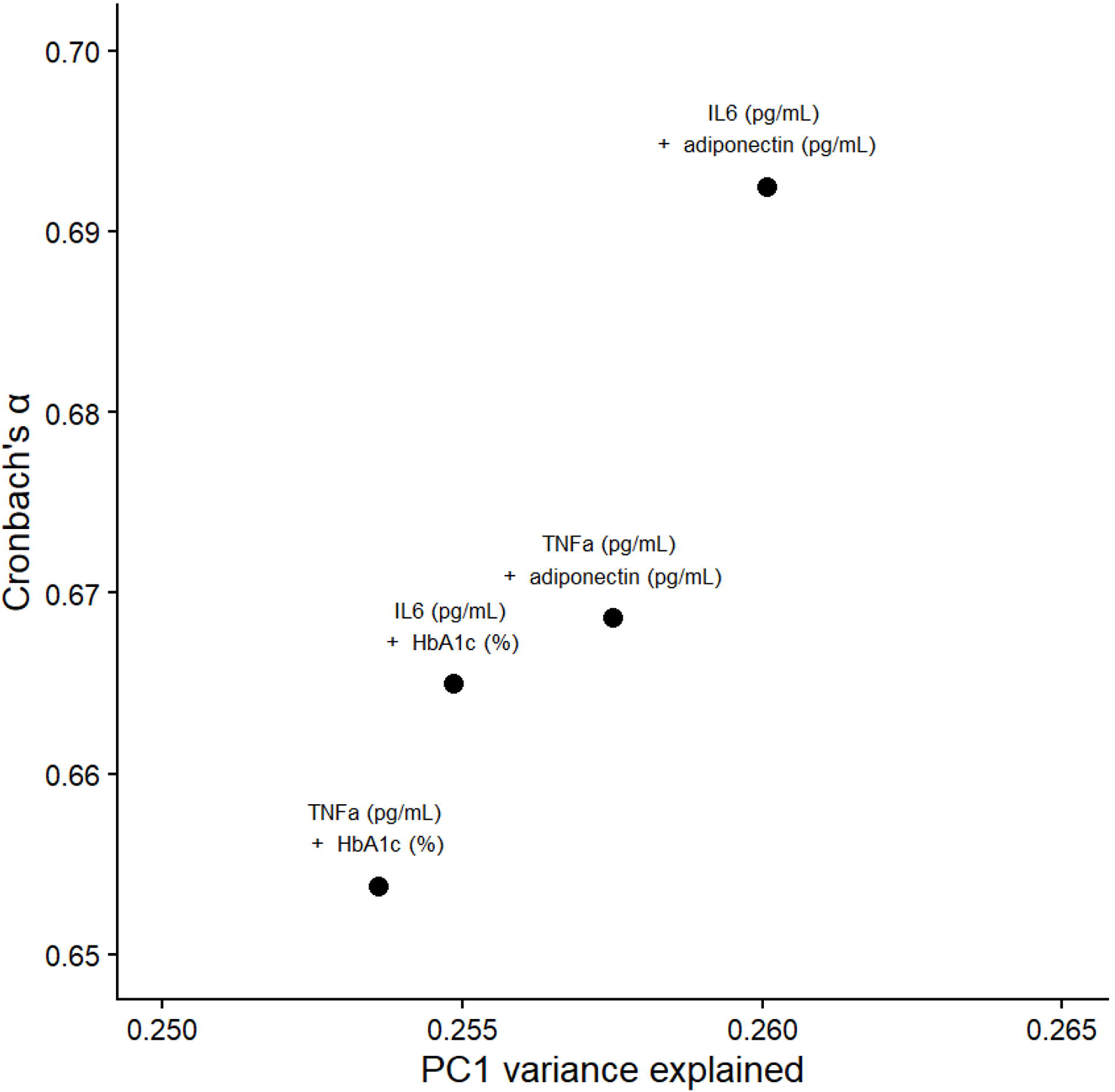

Domains-specific standardized scores were generated by calculating the mean standardized value within each domain to obtain a composite domain score (i.e., immune and stress response, energy and metabolism, and neuromuscular function). A composite VC score was subsequently derived by averaging the domain-specific Z-scores. Schematic visualisation of the VC score calculation is shown in figure 2.

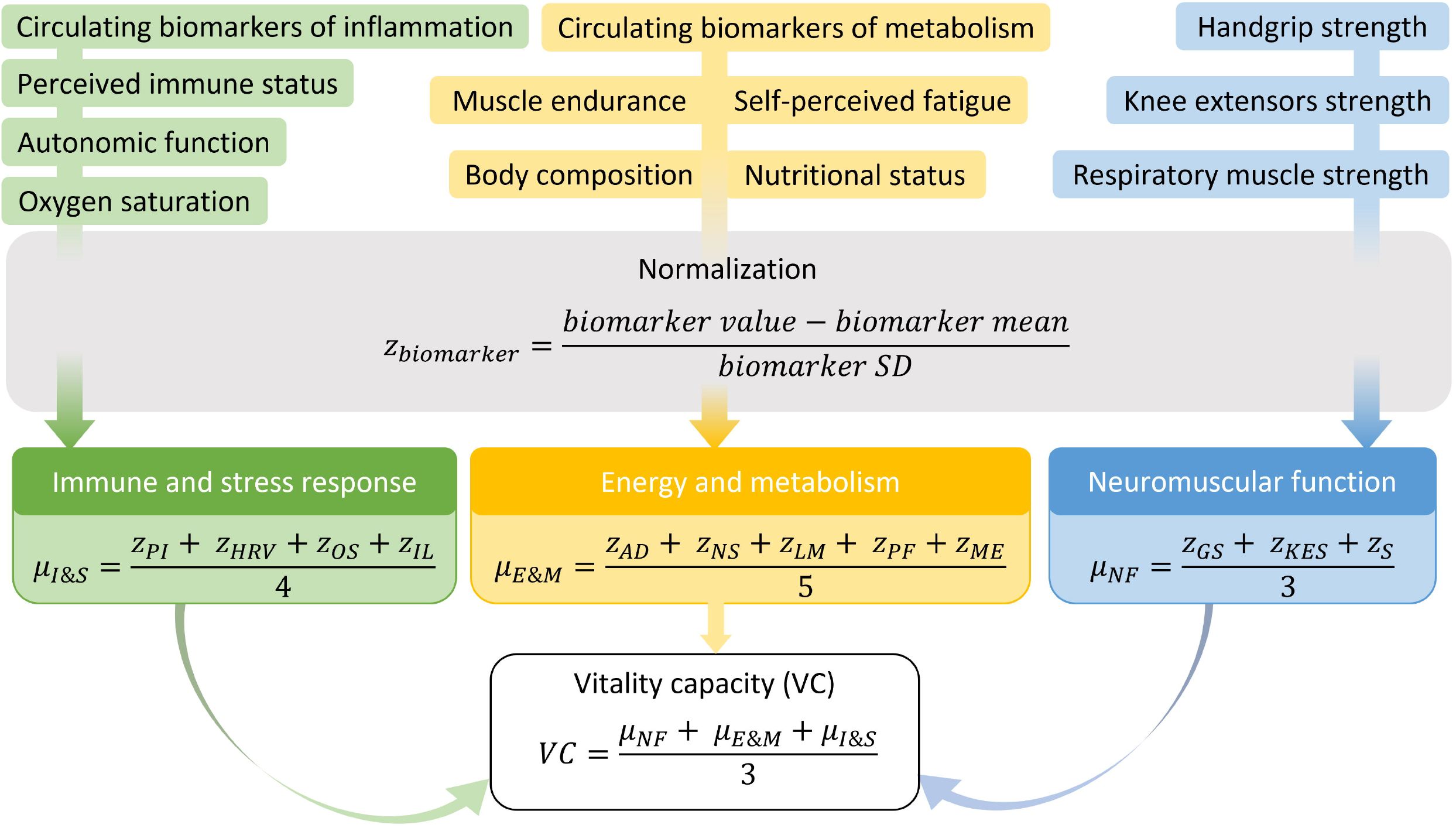

### 2.7. Statistical analysis

All statistical analysis were performed on R version 4.5.3 using the tidyverse R package version 2.0.0. Residual normality was assessed using the Shapiro–Wilk test and homoscedasticity using Levene’s test. Between-group comparisons were conducted using a 2-way ANCOVA including group and sex as fixed factors and age as a covariate. When a significant interaction effect was detected, post hoc pairwise comparisons were performed on estimated marginal means with p-values adjusted using the Benjamini–Hochberg procedure, performed using the emmeans package (version 2.0.2) in R. Results are presented as mean ± standard deviation, with effect sizes reported as partial eta-squared (η^2^p) for ANCOVA and Cohen’s d for post hoc comparisons.

Associations between the VC composite z-score and age, moderate to vigorous physical activity (MVPA) and quality of life, were assessed using Pearson correlations.

Principal component analysis (PCA) was performed on the VC domain scores. Participants were projected onto the first two principal components (PC1 and PC2). Group-level dispersion was illustrated using 95% confidence ellipses. Domain loadings were represented as vectors, indicating the direction and magnitude of their contribution to the principal components. The relative contribution of each domain to PC1 was calculated and expressed as a percentage of explained variance and visualized using a bar plot. Linear regression analysis and Pearson correlation were performed between PC1 scores and the VC composite z-score and visualized using a fitted linear regression line and 95% confidence interval.

A correlation heatmap was produced using the pheatmap R package (version 1.0.13) to provide an overview of the relationships between biomarkers, domains, and the VC composite score. Pearson or Spearman correlations were computed according normality distributions. The significance level was set at p< 0.05. All p-values were adjusted using Benjamini–Hochberg correction.

## 3. Results

### 3.1. Participants’ characteristics

Clinical and anthropometric characteristics of the study participants are shown in Table 1. A significant group × sex interaction was observed for body fat (F(1, 34) = 4.89, p = 0.034, η^2^p = 0.13). Active females had a lower body fat percentage compared to inactive females (t(34) =-4.204, p.adj < 0.001, d = -1.44).

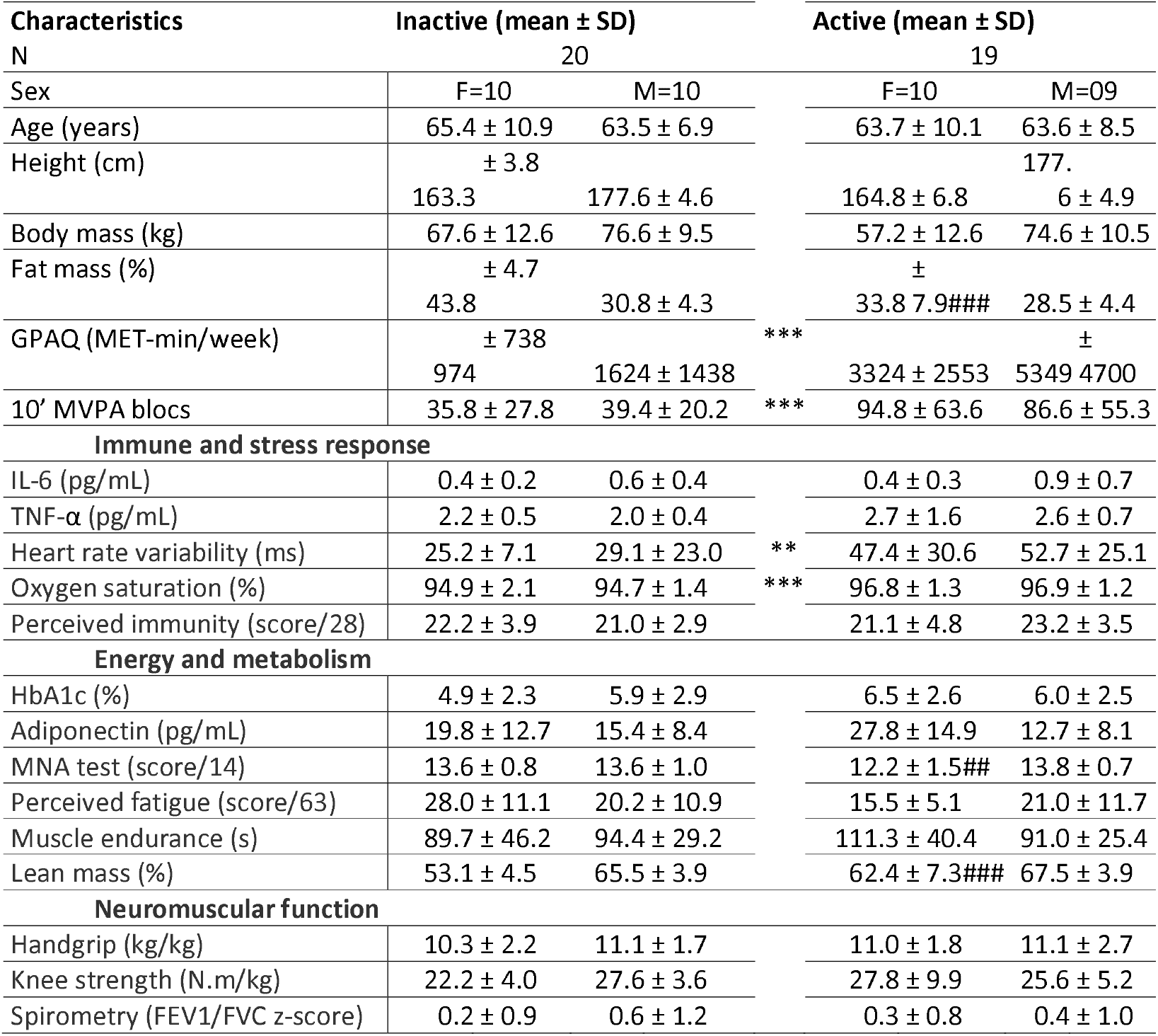

Significant group differences were observed for the GPAQ and accelerometry data (Table 1). Active participants exhibited significantly higher GPAQ scores (MET-min/week, 4283.2 ± 3762.9 vs 1299.0 ± 1161.5, F(1, 34) = 12.26, p = 0.001, η^2^p = 0.26) and time spent in > 10-min MVPA bouts (90.9 ± 58.3 vs 37.6 ± 23.7, F(1, 34) = 14.38, p < 0.001, η^2^p = 0.30), compared to inactive participants. No other significant difference was detected.

### 3.2. VC score

The composite VC score derived from averaging the domain scores was significantly higher in the active group compared with the inactive group (0.1 ± 0.3 vs −0.1 ± 0.4, F(1, 34) = 8.49, p= 0.006, η^2^p = 0.14). VC was inversely correlated with age (Fig. 3A, r(37) = −0.53, p < 0.001) and positively correlated with MVPA in the inactive group (Fig. 3B, ρ(n = 39) = 0.45, p = 0.048). No association was observed between VC and the quality-of-life score (Fig. 3C) nor with any WHOQOL-OLD subdimension.

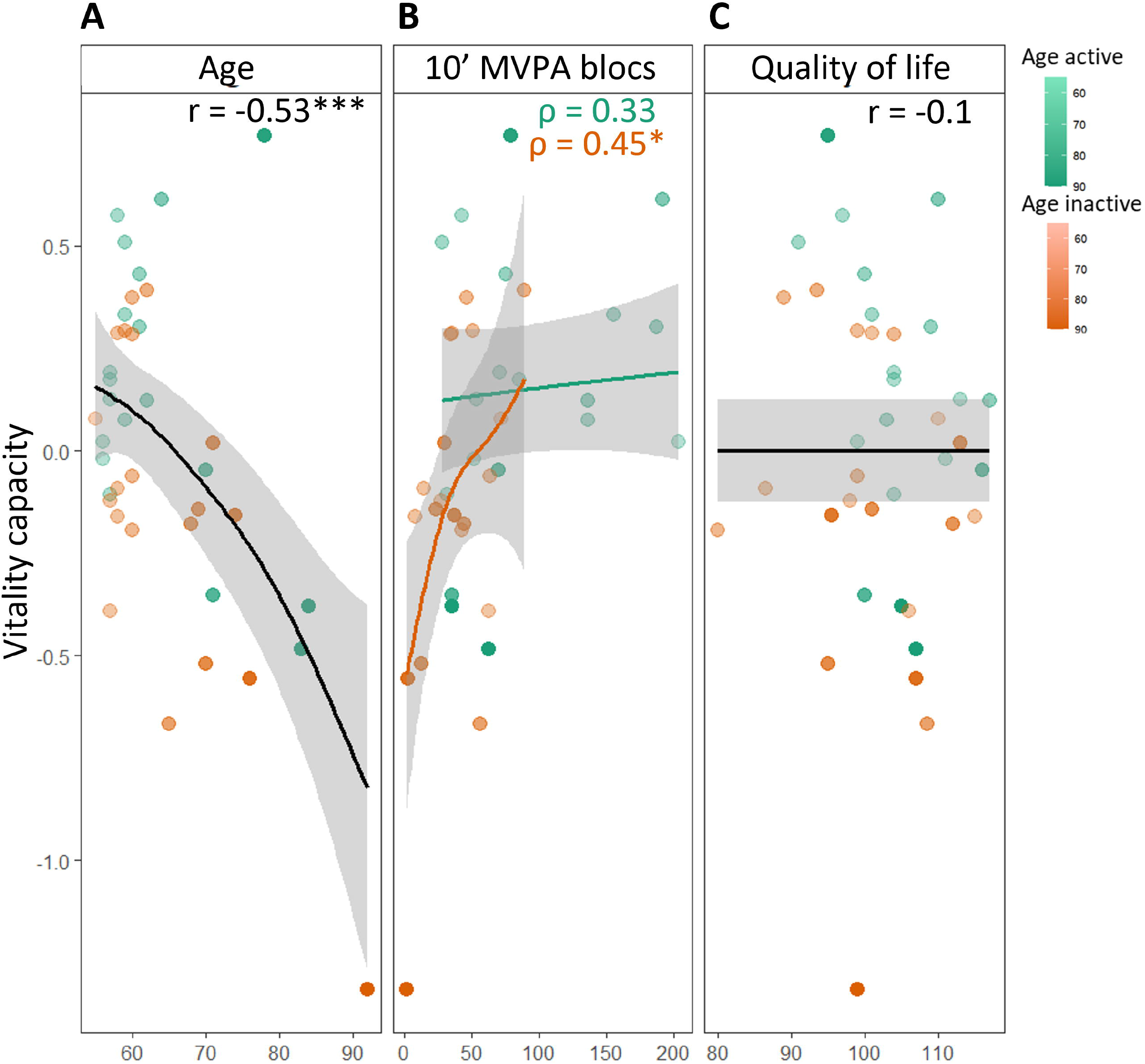

### 3.3. VC domains

A group effect was observed for immune and stress response score (F(1, 34) = 12.03, p = 0.001, η^2^p = 0.26). Active participants had a higher score than inactive participants (0.2 ± 0.5 vs -0.2 ± 0.5; Fig. 4A). A significant group × sex interaction effect was observed for energy and metabolism score (F(1, 34) = 4.14, p = 0.05, η^2^p = 0.11). After post-hoc analysis, only female active participants had a higher energy and metabolism score compared to inactive females (0.2 ± 0.3 vs -0.3 ± 0.4, t(34) = 2.950, p.adj = 0.006, d = 1.01). No significant main or interaction effects were observed in the neuromuscular function domain.

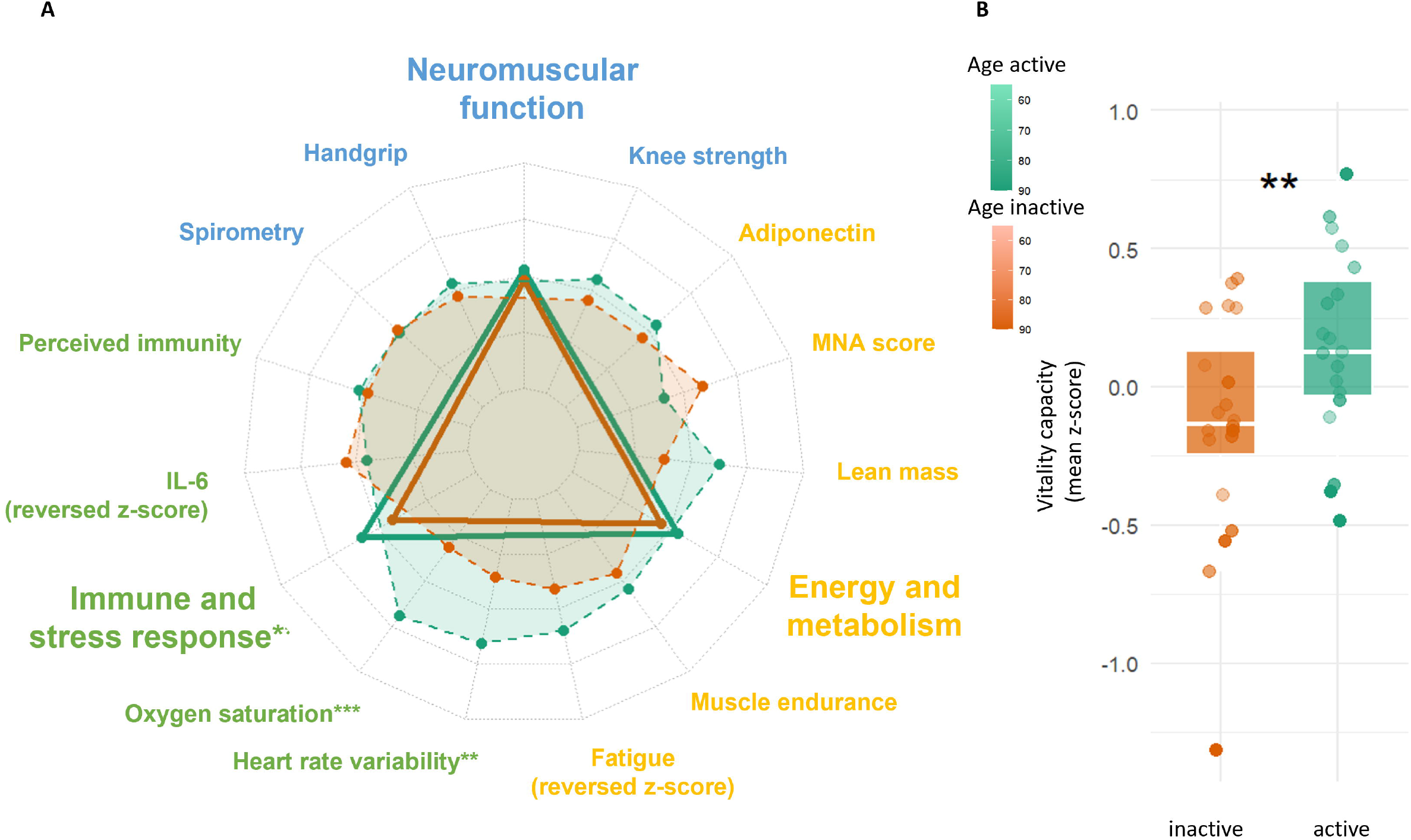

### 3.4. VC biomarkers

#### 3.4.1. Immune and stress response biomarkers

A group effect has been found for oxygen saturation and heart rate variability (RMSSD) (Table 1). Active participants had significantly higher oxygen saturation (%, 96.8 ± 1.2 vs 94.8 ± 1.8, F(1, 34) = 19.87, p < 0.001, η^2^p = 0.37) and heart rate variability (ms, 49.9 ± 27.5 vs 27.2 ± 16.7, F(1, 34) = 9.40, p = 0.004, η^2^p = 0.22) than inactive participants. No significant differences were observed in plasma IL-6 and TNF-α concentrations or perceived immunity (Table 1).

### 3.4.2. Energy and metabolism biomarkers

A significant group × sex interaction was found for lean mass (F(1, 34) = 5.28, p = 0.028, η^2^p = 0.13) and MNA test (F(1, 34) = 5.44, p = 0.026, η^2^p = 0.14). Female active participants had higher lean mass (t(34) = 4.232, p.adj < 0.001, d = 1.45) and lower MNA score (t(34) = -2.954, p.adj = 0.006, d = -1.01) compared to inactive females (Table 1). No significant differences were observed in blood HbA1c level, plasma adiponectin concentration, perceived fatigue or muscle endurance (Table 1).

#### 3.4.3. Neuromuscular function biomarkers

No significant effects were observed for neuromuscular biomarkers, including handgrip strength, knee strength, and FEV1/FVC z-score (Table 1).

### 3.5. Principal component analysis

PCA performed on VC domains identified a first principal component (PC1) that was highly correlated with the composite VC z-score (Fig. 5B). PCA of VC domains revealed that all domains contributed positively to PC1, although with different magnitudes (Fig. 5A). PC1 explained 62.2% of the variance and was mostly driven by neuromuscular function in both groups (Fig. 5A & 5C). In both groups, the neuromuscular function domain showed the strongest correlation with VC score (Fig. 5D, active, r(17) = 0.64, p.adj = 0.03, inactive r(18) = 0.83, p.adj < 0.001). Only the handgrip strength biomarker was significantly correlated with the VC score in the inactive group (Fig. 5D, r(18) = 0.76, p.adj = 0.003). In the active group, handgrip strength (Fig. 5D, r(17) = 0.84, p.adj < 0.001) and oxygen saturation (Fig. 5D, r(17) = 0.66, p.adj = 0.03) were correlated with VC.

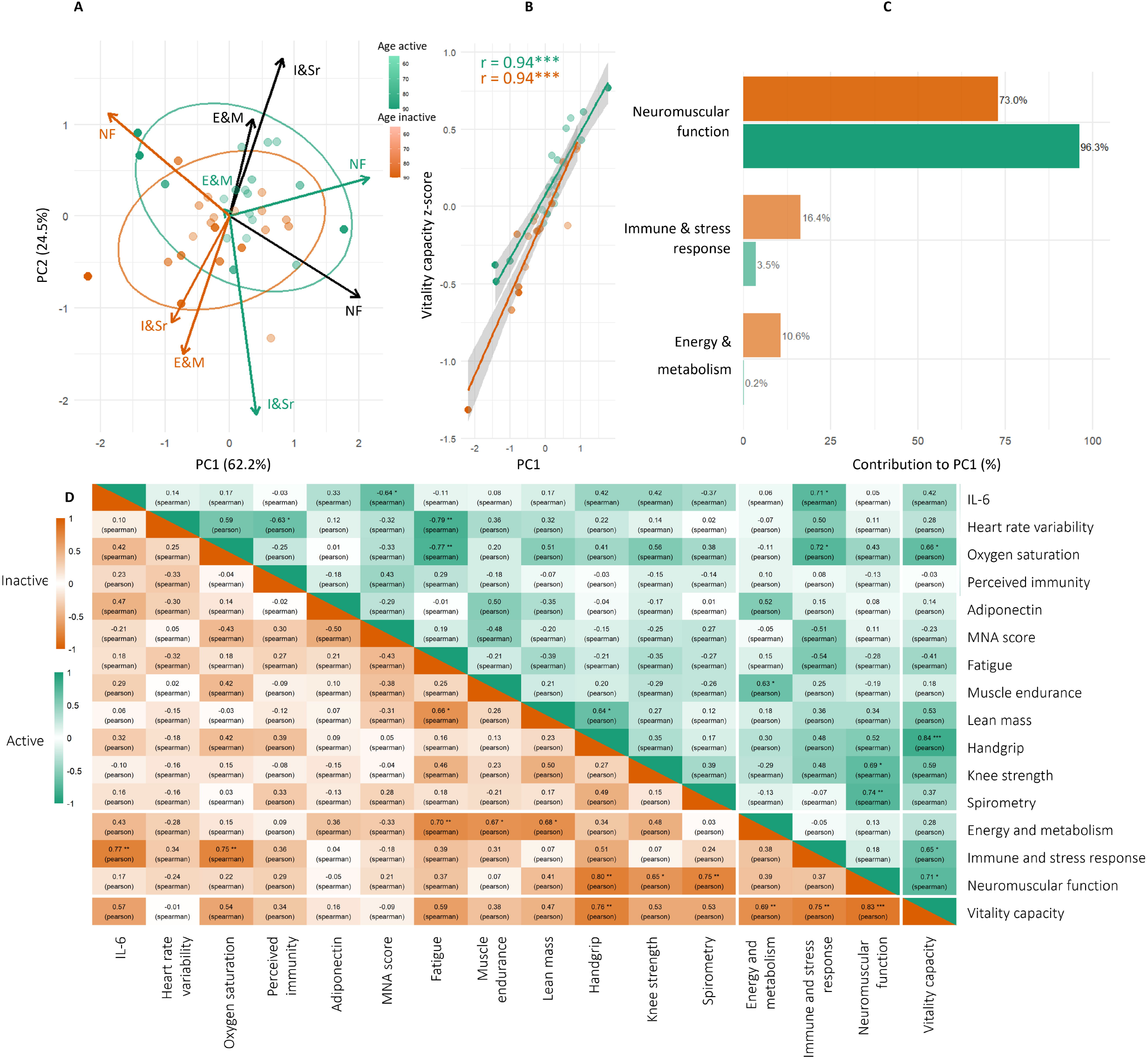

## 4. Discussion

This study aimed to measure and compare VC of lifelong aerobic active and inactive healthy older adults and estimate each domain weight in the VC construct. We report, for the first time, the operationalization of a VC score constructed from 12 biomarkers consistent with those proposed by the WHO working group on VC^14^. As hypothesized, lifelong aerobic active older adults had higher VC than inactive counterparts. We highlighted that, despite contributing substantially to overall VC variability, the neuromuscular function domain of VC did not significantly differ between groups. In contrast, the immune and stress response domain score, as along with some of its biomarkers (i.e., oxygen saturation, heart rate variability) showed the clearest distinction between lifelong aerobic active and inactive healthy older adults (Fig. 4A).

The inverse correlation between age and VC is consistent with the existing literature on VC^16^ and, more broadly, with the decline of intrinsic capacity with age^27^. The positive correlation between MVPA and VC, observed only in inactive (Fig. 3B), may suggest a plateau effect beyond a certain level of PA. This finding is consistent with large cohort studies and meta-analyses reporting non-linear relationships between levels of PA and intrinsic capacity^12^ and health related outcomes^28–30^. Even though quality of life has been positively associated with intrinsic capacity^31,32^, our study found no association between quality of life and VC among healthy older adults. This finding is consistent with a recent study^17^ investigating the relationship between VC and quality of life, which reported heterogeneous associations depending on the population studied and the assessment tools used. Overall, the absence of a clear association between quality of life and VC may reflect the complexity of the quality-of-life construct, which relies not only on individual physiology but also on a range of social and environmental factors.

Neuromuscular function showed the strongest contribution to the main principal component axis variability, which was highly correlated with VC, suggesting it might be a key contributor to inter-individual differences in VC. However, the biomarkers of neuromuscular function did not significantly differ between our lifelong aerobic active and inactive older adults. This contrasts with previous literature linking higher aerobic PA levels with greater handgrip and knee extension strength as well as spirometry measures^33,34^. One possible explanation is that the inactive group in our study consisted of relatively healthy individuals. Remaining healthy despite lower levels of PA may reflect, among other factors, a genetic predisposition to preserved functions. Several studies show that measures such as handgrip and knee extension strength, and spirometry lung function are heritable and are associated with specific genomic loci ^35–38^. Also, respiratory muscle function was assessed with spirometry, whereas a recent review suggested the use of volitional tests, which assess respiratory muscle strength^39^. Overall, although neuromuscular function appears to be a major domain of VC in our healthy population, the absence of between-group differences suggests that intrinsic factors, such as genetic predisposition may attenuate the impact of PA in relatively healthy older adults.

The immune and stress response domain also contributed to PC1 variance, although to a lesser extent. The higher immune and stress response domain score of lifelong aerobic active participants suggest that PA may improve it. Although scarcely investigated^18^, some biomarkers related to the immune and stress response domain showed the most pronounced differences between groups. The higher oxygen saturation observed in the active group is consistent with a recent study reporting a positive correlation between daily steps and oxygen saturation in older adults^40^. Similarly, the higher heart rate variability (HRV) in active older adults has been previously reported and supported by meta-analysis concluding that chronic endurance exercise improves HRV^41^. In contrast, plasma IL-6 concentrations did not significantly differ between groups, despite the well-established association between IL-6 and age-related diseases, as well as evidence suggesting a mitigating effect of lifelong exercise^7,42^. Perceived immunity did not differ between groups. The questionnaire used may not have been fully adapted to our healthy population; for instance, some reported symptoms (e.g., muscle pain^23^) may have reflected exercise-induced discomfort rather than illness. The limited scientific literature addressing this topic makes interpretation challenging and highlights the need for further investigations, potentially using a recently developed scale ^43^.

The energy and metabolism domain showed the lowest contribution to PC1 and no contribution in the active group. These findings suggest that this domain may be less discriminative among physically active older adults, possibly due to a more homogeneous metabolic profile within this group. Nevertheless, some biomarkers of the energy and metabolism domain significantly differed between females. Lifelong aerobically active females exhibited higher lean mass than their inactive counterparts, consistent with the well-established effects of PA on body composition ^44^. Because females typically have lower lean mass than males^45^, they may experience more pronounced physiological adaptations to PA, making the energy and metabolism domain more discriminating for females. The lower MNA scores observed in lifelong active females likely reflect their lower body mass index–commonly reported in females athletes^46^–rather than poorer nutritional status. This interpretation is consistent with a recent scoping review suggesting that MNA may not be fully adapted to healthy populations or individuals from high-income countries^18^. Collectively, these results suggest that sex-specific differences should be considered when interpreting VC, particularly for energy and metabolism domain. The absence of differences in muscle endurance, perceived fatigue and inflammation biomarkers (i.e., TNF-α and IL-6) is consistent with their known interrelationships in older adults^47^. Finally, adiponectin levels were within the normal range in our cohort^48^, and the lack of between-group differences is consistent with the similar HbA1c levels observed across groups, further reflecting the relatively healthy status of the study population.

The strong correlation of handgrip strength with VC reflects its association with different ageing trajectories^49–51^. In the active group, the significant correlation between oxygen saturation and VC, together with the strong association between lean mass and VC, supports the physiological relevance of the proposed VC construct and further strengthens the use of a physically active cohort as a relevant model of healthy ageing. These findings highlight the potential of a simplified biomarker-based approach, possibly based on handgrip strength, oxygen saturation, and lean mass. Further studies are needed to refine the selection of the most relevant biomarkers, which could enable a simplified VC assessment as an initial health-screening assessment.

This study has several strengths. First, it applies a multidimensional, biomarker-based approach to assess VC in accordance with the WHO framework. Second, the inclusion of lifelong aerobic active older adults provides a unique and physiologically relevant model of successful ageing. Finally, the limited sample size may also be considered a strength in the context of the growing emphasis on precision medicine. Indeed, the assessment of VC should be meaningful not only at the population level but also at the individual level, where inter-individual variability becomes critical. In this perspective, our approach supports the development of tools capable of capturing clinically relevant differences in physiological ageing trajectories. Some limitations should nevertheless be acknowledged. First, our biomarker selection required some methodological choices because the working group recommendations were sometimes broad^14^. Respiratory muscle strength was assessed using spirometry, which evaluate the respiratory muscle function, rather than maximal expiratory pressures^39^. The pair of blood biomarkers was not specified by the group^14^. We therefore selected 4 candidates based on literature and used an unbiased statistical approach to retain the pair that fitted the best in our population. The specificity (e.g., aged, Caucasians, healthy…) of our population may limit generalizability, and the absence of genetic data restricts the interpretation of inter-individual differences, particularly regarding neuromuscular function domain. Finally, although some biomarkers and assessment tools—particularly those related to blood, MNA and perceived immunity—may lack specificity in physically active populations, the overall selection remains robust. Considering that the proposed construct includes 12 biomarkers intended to be applicable across diverse populations, this represents a satisfactory compromise, highlighting the inherent challenge of identifying universally relevant biomarkers of VC.

## 5. Conclusion

In conclusion, our findings suggest that lifelong aerobic physical activity is associated with higher VC in older adults, supporting the concept that lifelong physically active individuals represent a relevant model for the study of healthy ageing. These results reinforce the World Health Organization framework of healthy ageing and highlight the potential of VC as an integrative measure of physiological ageing. Future longitudinal and experimental studies are needed to confirm these associations, refine biomarker selection, and establish reference values applicable at both individual and population levels. Expanding future research to more diverse cohorts will also be essential to improve generalizability across populations. Our findings further suggest that PA modalities may differently affect VC domains. While aerobic PA appears particularly relevant to improve the immune and stress domain, neuromuscular function may require complementary approaches such as resistance training. This, and the sex effect observed in the energy and metabolism domain, raises the possibility of more tailored PA programs according to individual physiological profiles. More broadly, this study emphasizes the fundamental role of sustained physical activity across the lifespan to preserve physiological functions, reinforcing the idea that maintaining movement is not only beneficial, but intrinsic to human healthy ageing.

## Acknowledgment

This work was supported by “Crédits Scientifiques Incitatifs” funding from Université Côte d’Azur and by the French government through the France 2030 investment plan managed by the National Research Agency (ANR), as part of the Initiative of Excellence Université Côte d’Azur under reference number ANR- 15-IDEX-01.

## Authors contribution

J. Mercier participated in the design of the study, contributed to data collection, analysis and interpretation of results, writing the article; S.S. Colson participated in the design of the study and interpretation of results, writing the article; A.S. Rousseau participated in the design and coordination of the study, contributed to data collection and interpretation of results, writing the article; F. Chorin contributed to data collection; E. Michel contributed to data collection; O. Guérin participated in the design of the study and project administration; N. Gautier contributed to data collection and analysis; A. Loubat contributed to data collection and analysis. All authors did read and approve the final version of the manuscript, and agreed with their position in the author list.

